# Exploiting long read sequencing to detect azole fungicide resistance mutations in *Pyrenophora teres* using unique molecular identifiers

**DOI:** 10.1101/2023.05.05.539008

**Authors:** Katherine G. Zulak, Lina Farfan-Caceres, Noel L. Knight, Francisco J. Lopez-Ruiz

## Abstract

Resistance to fungicides is a global challenge as target proteins under selection can evolve rapidly, reducing fungicide efficacy. To manage resistance, detection technologies must be fast and flexible enough to cope with a rapidly increasing number of mutations. The most important agricultural fungicides are azoles that target the ergosterol biosynthetic enzyme sterol 14α-demethylase (CYP51). Mutations associated with azole resistance in the *Cyp51* promoter and coding sequence can co-occur in the same allele at different positions and codons, increasing the complexity of resistance detection. Resistance mutations arise rapidly and cannot be detected using traditional amplification-based methods if they are not known. To capture the complexity of azole resistance in two net blotch pathogens of barley we used the Oxford Nanopore MinION to sequence the promoter and coding sequence of *Cyp51A*. This approach detected all currently known mutations from biologically complex samples increasing the simplicity of resistance detection as multiple alleles can be profiled in a single assay. With the mobility and decreasing cost of long read sequencing, we demonstrate this approach is broadly applicable for characterizing resistance within known agrochemical target sites.

## Introduction

Fungicides are an important tool for the management of crop disease with the use of demethylase inhibitor or azole fungicides dominating in both agriculture and clinics (Kelly *et al*., 1993, Parker *et al*., 2014). Azoles are single site systemic fungicides that target the cytochrome P450 sterol 14α-demethylase (CYP51) enzyme, which has an essential role in the biosynthesis of the fungal sterol ergosterol (Lamb *et al*., 1999). Azole fungicides disrupt the function of CYP51 and therefore membrane stability and fluidity through a combination of ergosterol depletion and the accumulation of toxic sterol intermediates (Cowen *et al*., 2014).

Azole fungicides are highly efficient and resistance development is generally slower than other single site fungicides with incomplete cross resistance among members of the class (Parker et al., 2014, Cools *et al*., 2013). However an increasing number of cases of azole resistance have been reported among economically important crop pathogens such as *Zymoseptoria tritici* (Cools & Fraaije, 2013), *Fusarium graminearum* (Becher *et al*., 2011), *Blumeria graminis* spp. (Tucker *et al*., 2015, Wyand & Brown, 2005), and *Pyrenophora teres* (Mair *et al*., 2016, Mair *et al*., 2020); as well as human fungal pathogens such as *Aspergillus fumigatus* (Chamilos & Kontoyiannis, 2005). There are three main mechanisms of azole fungicide resistance; mutations in the *Cyp51* gene that alter fungicide binding, over-expression of *Cyp51* due to promoter modifications, and increased efflux out of the cell (Cools et al., 2013). These mechanisms are not mutually exclusive, and several can exist in the same isolate, increasing resistance levels. A further complication is the existence of more than one paralogous *Cyp51* gene in the filamentous Ascomycetes, including many phytopathogens. However, in ascomycetes with more than one *Cyp51* paralogue, mutations associated with fungicide resistance are associated with only *Cyp51A* (Mair et al., 2016, Fan *et al*., 2013, Brunner *et al*., 2015).

*P. teres* f. *teres* (*Ptt*) and *P*. *teres* f. *maculata* (*Ptm*) are the causal agents of net form of net blotch and spot form of net blotch of barley (*Hordeum vulgare*) respectively. *Ptt* contains one *Cyp51B* gene and two copies of *Cyp51A*; *Cyp51A1* and *Cyp51A2*, with the fungicide resistance mutation F489L only occurring in *Cyp51A1* (Mair et al., 2016). Expression of all three genes was induced by tebuconazole but no promoter changes were found (Mair et al., 2016). Unlike *Ptt*, *Ptm* contains only one copy of *Cyp51A*, but both promoter and coding sequence mutations are associated with azole fungicide resistance (Mair et al., 2020). Solo-long terminal repeat (LTR) insertion elements were present in the *Cyp51A* promoter at five different locations and the F489L mutation was encoded by three different non-synonymous mutations (Mair et al., 2020). Isolates containing both insertions and the F489L mutation showed high resistance to azoles, while isolates containing either insertions or F489L had a moderately resistant phenotype (Mair et al., 2020). *Ptt* and *Ptm* can also form hybrids in nature (Campbell *et al*., 2002, Lehmensiek *et al*., 2010, Leišova *et al*., 2005, McLean *et al*., 2014, Turo *et al*., 2021) with one hybrid haplotype exhibiting a high level of resistance to azole fungicides suggested to have acquired the trait through natural recombination between the two forms (Turo et al., 2021, Mair et al., 2020).

Detection and monitoring of fungicide resistance in both agriculture and clinics is critical for monitoring resistant populations and informing appropriate management. For example, if resistance to an azole compound is detected, chemical management can be adjusted to avoid placing excessive selective pressure on the pathogen population and then the population closely monitored to ensure the treatment change was effective. This requires rapid, accurate and point of care molecular detection technologies capable of capturing the complex network of both known and evolving genetic mutations that cause resistance. Several allele-specific technologies such as loop-mediated isothermal amplification, digital PCR, and quantitative PCR have been deployed for the detection and monitoring of fungicide resistance in phytopathogens (Fan *et al*., 2018, Duan *et al*., 2014, Zulak *et al*., 2018, Hellin *et al*., 2021, Dodhia *et al*., 2021, Knight *et al*., 2023). Although useful, these technologies suffer from two major limitations, the need for a separate assay for each mutation and the inability to detect new mutations. In *Ptt* and *Ptm* there are several different promoter insertions and coding sequence mutations that can exist separately or in combination (Mair et al., 2016, Mair et al., 2020). *Ptm* and *Ptt* are also closely related and can form hybrids (Turo et al., 2021, McLean et al., 2014), further complicating resistance diagnoses. Thus, screening for resistance from field samples becomes a complicated and time-consuming process involving a large array of assays. In addition, the F489L mutation is encoded by three codons in *Ptm* (Mair et al., 2020) which could result in a false negative if these changes were not known.

Amplicon sequencing of marker genes such as 16S ribosomal DNA for prokaryotes and 18S rDNA and internal transcribed spacers (ITS) for eukaryotes has been used in the field of metagenomics to characterize microbial diversity, mainly using high throughput short read sequencing technologies such as Illumina (Liu *et al*., 2021). However, these short read lengths can constrain taxonomic resolution to family or genus level and even confuse taxonomic assignment if different parts of the marker gene are used (Schloss, 2010). Longer read sequencing technologies such as Pacific Biosciences (PacBio) and Oxford Nanopore Technologies (ONT) could remedy this issue but suffer from relatively high error rates and resolving single nucleotide polymorphisms is problematic. PacBio has resolved this issue through circular consensus sequencing to generate high accuracy (99.8%) long reads (Wenger *et al*., 2019). Recently, this was exploited to produce error-corrected full length rRNA genes and resolve the majority of a bacterial mock community to species level (Earl *et al*., 2018). This technology was also used recently to characterize azole, succinate dehydrogenase inhibitor (SDHI), and quinone outside inhibitor (QoI) resistance in the phytopathogen *Z. tritici* (Samils *et al*., 2021). Target genes were multiplexed, amplified, and sequenced using PacBio circular consensus sequencing to characterize the range of resistance genotypes present in European populations of *Z*. *tritici*. The authors also included nine housekeeping genes to further profile the strains present in these populations (Samils et al., 2021). This study presents a comprehensive system capable of profiling target genes to three modes of action, however samples must be sent to centralized laboratories to perform sequencing and this method involves culturing individual isolates, both of which can be time consuming.

ONT employs long read technology with the added benefit of portability which enables sequencing on demand that can be done in any laboratory or in the field, an attractive prospect for rapid resistance diagnosis. Recently, ONT sequencing was used to profile azole resistant haplotypes in the phytopathogen *Z. tritici* (Gutierrez Vazquez *et al*., 2022). The authors sequenced mock communities with varying quantities of one to three isolates as well as infected leaves to identify haplotypes associated with azole resistance. To correct sequencing errors, reads were filtered using Filtlong (https://github.com/rrwick/Filtlong) and corrected using Canu (Koren *et al*., 2017), which resulted in a per position consensus between 98 and 99%, although the filtering step resulted in substantial data loss and the potential effect of PCR chimeras was not discussed. Several additional pipelines have been developed to mitigate the per base 2 to 25% error rate (Wick *et al*., 2018) such as denoising (Kumar *et al*., 2019, Hathaway *et al*., 2018) and intramolecular-ligated nanopore consensus sequencing (Calus *et al*., 2018, Li *et al*., 2016), and recently unique molecular identifiers (UMIs) which tag individual molecules with a UMI sequence containing random bases in a particular pattern which can be used to bioinformatically sort molecules based on their original templates (Karst *et al*., 2021). This method results in a low consensus error rate of 0.0042% and a chimera rate of less than 0.02% when sequenced on an R10.3 flow cell (Karst et al., 2021). Chimeras are artifacts which arise when two or more sequences are joined together incorrectly and then further amplified and often occur in complex mixed samples. These artifacts are problematic because they are needlessly sequenced and can affect the perception of the DNA pool. In fact, chimeras can be upwards of 20% of the PCR products in metagenomics studies, depending on polymerase and cycling conditions (Marc & Patrick, 2019). The high accuracy and low chimera rate of UMIs make this method ideally suited to single nucleotide polymorphism (SNP)-level detection of both known and new mutations in fungicide target genes in complex biological samples with highly similar sequences.

Here we applied the UMI amplicon sequencing approach to profile both inserts and single nucleotide changes in the azole target gene *Cyp51A* in two forms of *P. teres* and a hybrid. We validated this approach using a mix of currently known haplotypes of *Ptt* and *Ptm* and resolved all variants with 100% identity to reference sequences. We then optimized a workflow to process diseased leaf samples and profiled their *Cyp51A* haplotypes and validated this using quantitative PCR, resulting in the detection of a point mutation associated with azole resistance in *Ptt*. This method is broadly applicable and can be used not only to profile fungicide target gene mutations, but also to characterize agrochemical resistance in other kingdoms such as bacteria, plants and insects.

## Materials and Methods

### Isolate selection and culturing

Seven *P. teres* isolates used in this study are listed in Table 1 and Figure 1. The details of all isolates have been previously published (Mair et al., 2016, Mair et al., 2020). A single conidium was isolated, grown and stored at −80°C as in Mair et al. (2016). Mycelial plugs were then grown on V8 potato-dextrose agar (PDA; 10 g potato-dextrose agar, 3 g CaCO3, 15 g agar, 150 mL V8 juice in 850 mL sterile deionized H_2_O) plates and incubated at room temperature in the dark for five to seven days.

**Figure 1.**
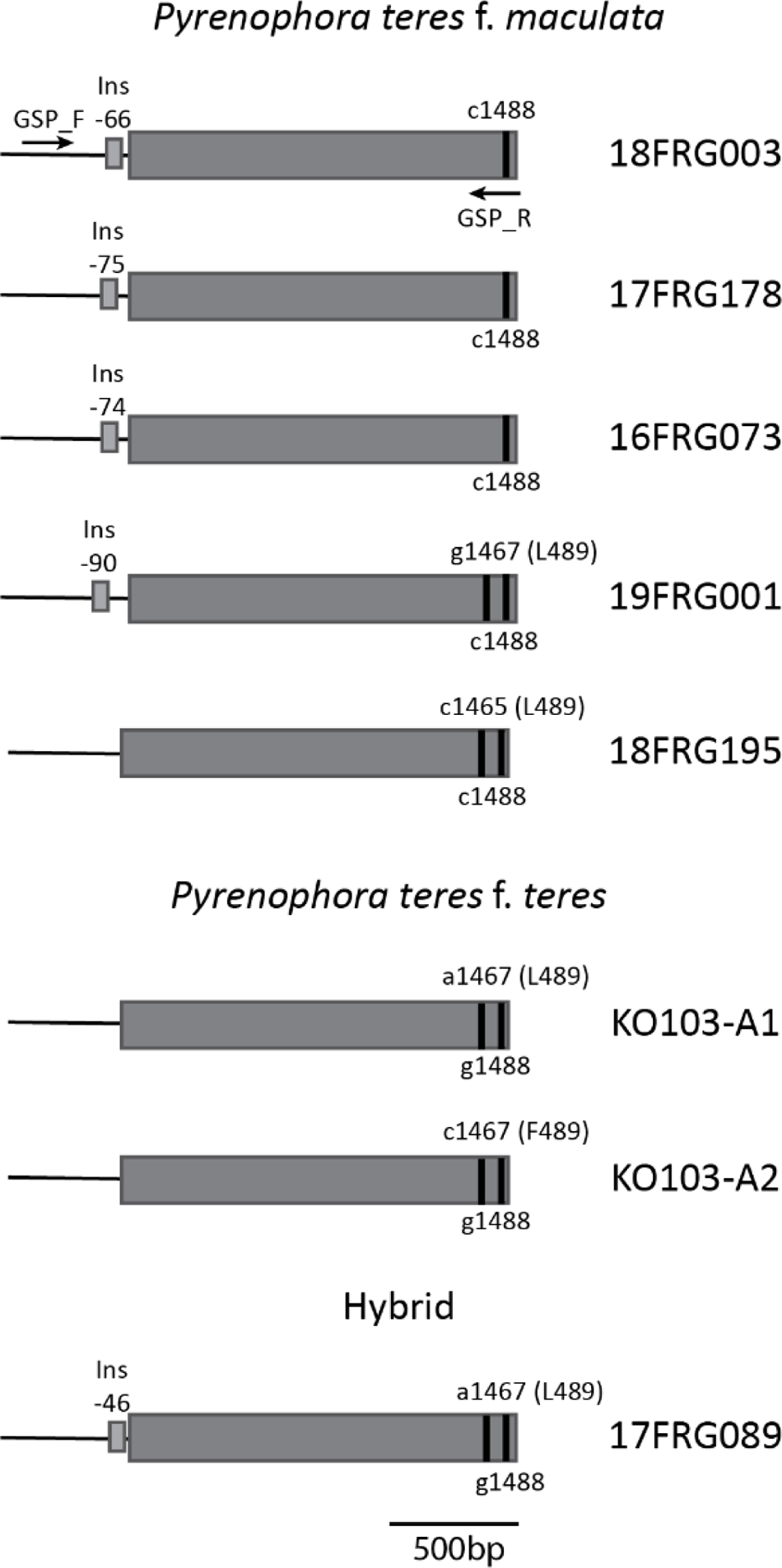
Genotypes of the azole fungicide target gene *Cyp51* in seven isolates of *Pyrenophora teres* f. *maculata*, *P. teres* f. *teres* and a hybrid amplified and sequenced in this study. Arrows indicate approximate primer positions and vertical black lines indicate nucleic acid positions associated with *P. teres* form differentiation and mutations associated with azole fungicide resistance (L489) and sensitivity (F489). Ins = promoter insertion associated with azole fungicide resistance.

**Table 1.** *Ptt* = *Pyrenophora teres* f. *teres*, *Ptm* = *Pyrenophora teres* f. *maculata*, MR = moderately resistant, HR = highly resistant, CDS = coding sequence

| Isolate ID | Year | Form | Location | Resistance phenotype | Reference | Genbank accession promoter | Genbank accession CDS |
| --- | --- | --- | --- | --- | --- | --- | --- |
| KO103 | 2013 | *Ptt* | Kojonup, WA | MR | Mair et al., 2016 |  | KX578217.1 |
| 16FRG073 | 2016 | *Ptm* | Gibson, WA | MR | Mair et al., 2020 | MT499783.1 | ON534218.1 |
| 17FRG178 | 2017 | *Ptm* | Coomalbidgup, WA | MR | Mair et al., 2020 | MT499784.1 | MT499777.1 |
| 18FRG003 | 2018 | *Ptm* | Munglinup, WA | MR | Mair et al., 2020 | MT499785.1 | ON534219.1 |
| 18FRG195 | 2019 | *Ptm* | Pithara, WA | MR | Mair et al., 2020 |  | MT499777.1 |
| 19FRG001 | 2019 | *Ptm* | Broomehill, WA | HR | Mair et al., 2020 | MT499787.1 | MT499780.1 |
| 17FRG089 | 2017 | *Ptm/Ptt* hybrid | Takaralup, WA | HR | Turo et al., 2021<br>Mair et al., 2020 | MT499786.1 | MT499779.1 |

### Primer design

*Cyp51A* promoter and coding sequences from *Ptt*, *Ptm*, and hybrid isolates used in this study were aligned with the *Ptt* and *Ptm* reference isolates W1 and SG1, respectively (GenBank accessions for seven isolates listed in Table 1; W1 = KX578221.1 and KX578220.1; SG1 = MT499781.1 and MT499776.1). Conserved regions in the promoter and 3′ sequence end were manually identified that captured all known promoter and coding sequence mutations. The forward gene specific primer is positions 302 to 320 bp in the SG1 reference MT499781.1 and the reverse gene specific primer is positions 1495 to1512 bp in the SG1 reference MT499776.1. Primers (Macrogen, Seoul) were tested by running PCR and analysing fragments through gel electrophoresis using genomic DNA from the seven isolates to ensure only one amplicon was present at the correct size. Universal primers and UMI tagging sequences were added to the gene specific primers as in Karst et al. (2021). All primers are listed in Table 2.

**Table 2.**
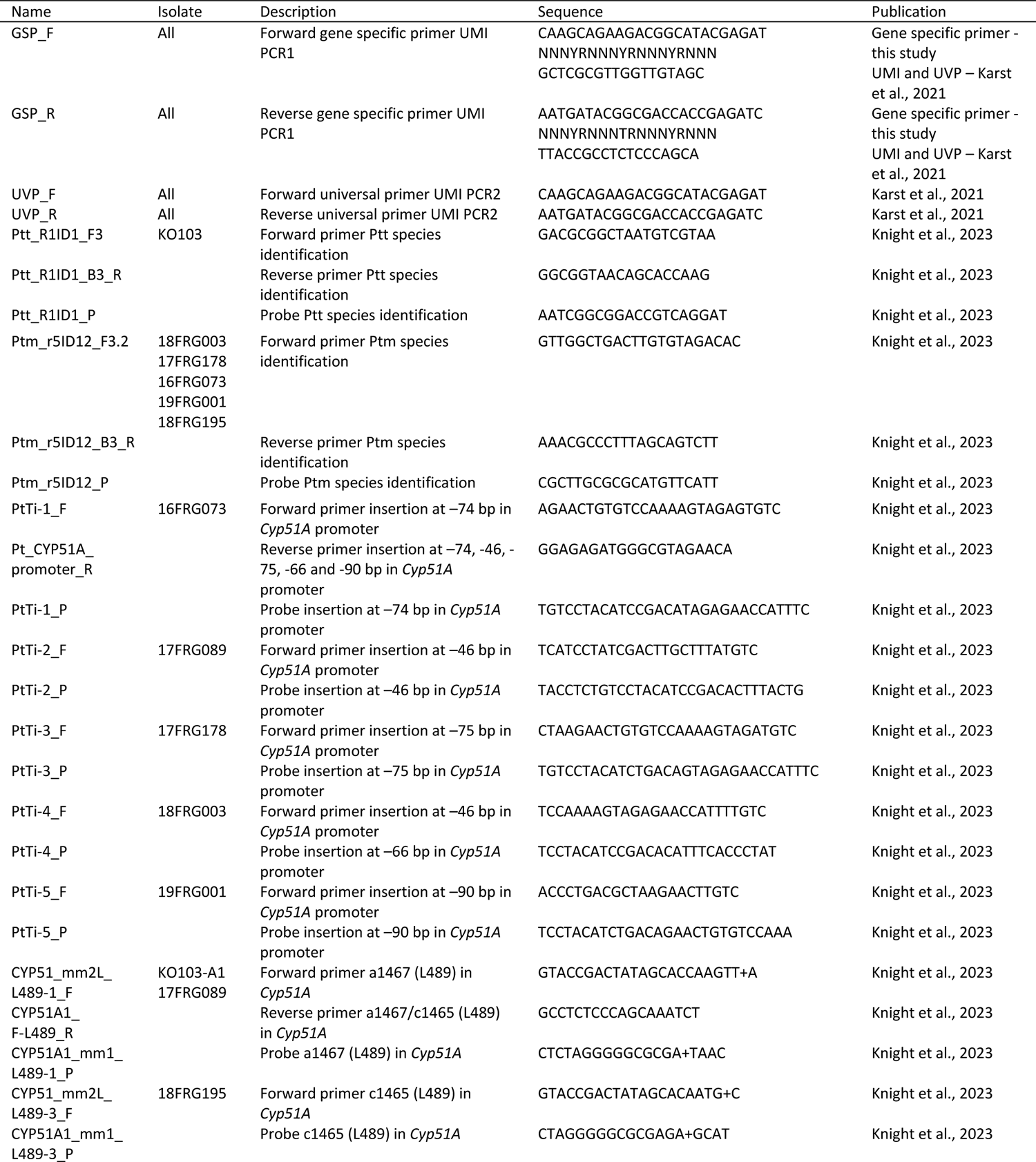
Primers used in this study.

### DNA extraction and UMI PCR of *Cyp51A* mock community

To create the *Cyp51A* mock community, DNA was extracted and quantified separately from each of seven *P. teres* isolates and then mixed in equimolar amounts before UMI PCR was performed. UMI PCR, library preparation and sequencing were then performed on the mixed sample. Mycelia from seven *P. teres* isolates was harvested using a scalpel blade from the culture plate, and separately placed in 1.5 mL tubes with two ball bearings and flash frozen in liquid nitrogen. Samples were then ground to a fine powder in a high-speed mixer mill (Retsch MM301, Germany) at 30 rev s^-1^ for one minute. DNA extractions were carried out using the Biosprint 15 DNA Plant Kit and Biosprint 15 robot (Qiagen, Australia) according to the manufacturer’s instructions. DNA concentrations were determined on a Qubit Flex Fluorometer using a Qubit dsDNA HS assay kit (Thermo Fisher). Optimal input DNA concentrations for ONT sequencing were calculated to be 1.3ng based on the 50 000 to 60 000 target strand copies recommended in a similar protocol published by ONT called “Custom PCR UMI” found at https://community.nanoporetech.com/docs/prepare/library_prep_protocols/custom-pcr-umi/v/cpu_9107_v109_revc_09oct2020?devices=minion.

Amplicons tagged with UMIs were generated following Karst et al., 2021 and using gene specific and universal primers listed in Table 2. First, a 2117 to 2243 bp region of the *Cyp51A* promoter and coding sequence (Figure 1) was amplified and tagged with UMIs using GSP_F and GSP_R primers (Table 2). The tagging PCR reaction contained 0.5 µM of each primer, 12.5 µL of 2X Platinum SuperFi II PCR Master Mix (0.08 U µL^-1^,Thermo Fisher Scientific), 1.3 ng of gDNA from each isolate (total of 9.1 ng genomic DNA; Table 1) for a total reaction volume of 25 µL. PCR conditions were as follows: initial denaturation at 95°C for 3 min followed by 2 cycles of 95°C for 30 s, annealing at 60°C for 30 s, followed by final extension at 72°C for 6 min. The PCR products were purified using AMPure XP beads (Beckman Coulter, California, USA) following manufacturer’s instructions except for a 0.6 bead to sample ratio and 80% ethanol concentration. Cleaned PCR products were then dissolved in 10 µL of ultrapure water.

A second PCR was performed using UVP_F and UVP_R (Table 2) to further amplify the UMI-tagged PCR products. The reaction contained 0.5 µM of each primer, 25 µL of 2X Platinum SuperFi II PCR Master Mix and the cleaned UMI-tagging reaction previously described, for a total reaction volume of 50 µL. PCR cycling conditions were as follows: initial denaturation 95°C for 3 min, followed by 25 cycles of 95°C for 30 s, annealing at 60°C for 30 s, 72°C for 6 min, followed by final extension at 72°C for 5 min. The resulting PCR products were cleaned using AMPure XP beads (Beckman Coulter, California USA). The purification was performed as described above except for 0.9 bead to sample ratio was used. PCR products were then resuspended in 10 µL of ultrapure water

To achieve the required number of amplicons for sequencing, a third PCR was carried out with the 20 ng of cleaned PCR products from the second PCR as template,0.5 µM of primers UVP_F and UVP_R and 50 µL of 2X Platinum SuperFi II PCR Master Mix, in a 100 µL reaction and 11 cycles at the conditions described for the second PCR. PCR products were purified as above for the second PCR and final amplicon concentration measured using a Qubit Flex Fluorometer and Qubit dsDNA HS assay kit (Thermo Fisher Scientific).

### Quantitative PCR

Quantitative PCR was performed to validate both the mock community and field samples using a magnetic induction cycler (MIC) qPCR system (Bio Molecular Systems, Australia) with two technical replicates per sample. Positive controls consisted of genomic DNA from the reference isolate known to contain the mutation being tested. Negative controls consisted of genomic DNA from an isolate known not to contain the mutation being tested. Each reaction mixture consisted of 10 uL of ImmoMix™ (Meridian Life Science, USA), 0.25µM of forward and reverse primers, 0.15 µM of probe and 2.5 ng of template gDNA for a total reaction volume of 20 µL. Assays were developed by Knight et al. (2023) and all primers and probes used are listed in Table 2. The cycling conditions were as follows: initial denaturation of 95°C for 10 min followed by 35 cycles of 95°C for 15 s and 62°C for 60s. Amplification and melt curves were analysed using the micPCR software version 2.10.0 (Bio Molecular Systems). Thresholds were automatically determined using the software and the quantification cycle (Cq) value was determined. A Cq value of less than 35 cycles was considered confirmation of the presence of a particular mutation.

### ONT sequencing of *Cyp51A* mock community

Library preparation was carried out using the amplicons from the third PCR reaction and the ONT Ligation Sequencing Kit (SQK-LSK109) following the ‘Amplicons by Ligation’ protocol as per instructions except for an initial amplicon concentration of 400 fmol. Sequencing was performed on a R10.3 MinION flow cell (FLO-MIN111) and monitored using MinKnow software version 21.02.5. Every 24 hours the flow cell was flushed using the ONT Flushing Buffer as per manufacturer instructions and another aliquot of library was loaded to obtain the maximum amount of data. Basecalling was performed using Guppy version 5.0.7 with a minimum q-score of 7. Mutations were validated using qPCR as above.

### Culturing of field samples

Net-blotch symptomatic leaf samples collected in 2021 from barley crops in Minlaton, South Australia, were assessed for fungicide resistance following a culturing method described by Knight et al. (2023). Briefly, infected barley leaves were surface sterilised for 30 s in 70% (v/v) ethanol, 60 s in 0.125% (w/v) NaOCl and 2× 60 s in autoclaved filtered water, lesions excised, and placed on 2 mL PDA supplemented with streptomycin (10µg mL^-1^), ampicillin (100µg mL^-1^) and neomycin (50µg mL^-1^) dispensed into a 12 well plate. Plates were incubated for seven days at 21°C in the dark. Resulting fungal growth was scraped from the media and macerated using ball bearings and a mixer mill (Retsch MM400). The mycelial fragments were then distributed onto 24-well plates containing 1 mL PDA with antibiotics as above and either no fungicide, tebuconazole at 15 µg mL^-1^ and 50 µg mL^-1^, and then incubated at 21°C in the dark. Fungal growth from each leaf sample was visually assessed after seven days and hyphae growing at both tebuconazole concentrations were harvested.

### ONT sequencing of field samples

Genomic DNA of mycelium harvested from both the 15 µg mL^-1^ and 50 µg mL^-1^ tebuconazole wells for each lesion (one lesion per well) were pooled in equimolar amounts and 1.3 ng of each mix was used as template for UMI PCR as described above. Library preparation was carried out using the ONT Ligation Sequencing Kit (SQK-LSK109) following the ‘Native barcoding amplicons’ protocol as per instructions except for an initial amplicon concentration of 500 fmol. The five samples were barcoded and sequencing was performed on a R10.3 MinION flow cell (FLO-MIN111) and flushed every 24 hours as described above. Basecalling was performed using Guppy version 5.0.12 with a minimum q-score of 7. Mutations were validated using qPCR as above.

### Data analysis

Raw passed fastq reads from the five samples were combined and analysed using the longread UMI nanopore pipeline (Karst et al., 2021) with the following settings: check start of read: 200, check end of read: 200, minimum read length: 2000, maximum read length: 5000, forward adaptor sequence: CAAGCAGAAGACGGCATACGAGAT, forward primer sequence: GCTCGCGTTGGTTGTAGC, reverse adaptor sequence: AATGATACGGCGACCACCGAGATC, reverse adaptor primer: TTACCGCCTCTCCCAGCA, racon consensus rounds: 2, medaka consensus rounds: 2, medaka model: r103_min_high_g345, bin size cutoff: 10. Consensus sequences from the variants.fa output file were aligned against reference *Cyp51A* sequences using Geneious Prime software version 2021.1.1 (Biomatters Ltd, Auckland New Zealand).

## Results

### UMI PCR assay design and sequencing optimisation

To ensure our assay detected all known *Cyp51A* variants to date, five *Ptm,* one *Ptt* and one *Ptm/Ptt* hybrid isolate that represent all currently known *Cyp51A P. teres* haplotypes were selected for amplicon sequencing (Figure 1, Table 1). We aligned the promoter and coding sequences of the seven isolates and designed gene-specific primers to capture all known promoter inserts, the F489L mutations and the g1448c SNP that discriminates *Ptt* from *Ptm* and the hybrid (Figure 1, Table 2). This resulted in an amplicon that captured all known mutations related with azole resistance in *P. teres*. Primers were validated on genomic DNA from individual isolates and resulted in bands of the expected sizes (18FRG003, 17FRG178, 16FRG073, 19FRG001 and 17FRG089 = 2327 bp; 18FRG195 = 2193 bp; KO103-A1 = 2191 bp; KO103-A2 = 2201 bp).

For the *Cyp51A* mock community sequencing, we determined the optimal amount of input DNA to be 1.3ng by empirically testing sequencing runs using 10, 5, 3 and 1.3 ng of gDNA from each of the seven isolates pooled into one PCR reaction (Table 1) and found the pipeline yielded sufficiently high cluster densities with 1.3 ng of gDNA from each isolate but not with the higher starting concentrations. PCR reactions using less than 1.3 ng template DNA per isolate failed to yield product.

### ONT sequencing of mock community resolves all variants with 100% accuracy

We sequenced the amplicons from the seven isolate mock community using the MinION, which resulted in 2,164,642 passed sequences. To determine the minimum number of reads per target for cluster formation and variant calling, we inputted a randomly sub-sampled set of 1.5M, 1M, 750K, 500K and 250K reads from the original dataset into the longread UMI pipeline (Figure 2, Table 3). The pipeline failed to generate clusters using 250K reads, but generated clusters and called variants using ≥500K sequences (Figure 2, Table 3). The 500K read input failed to identify three haplotypes, whereas the ≥750K read inputs resolved all haplotypes (Figure 2, Table 3). Although equimolar amounts of each isolate were used as input into the UMI PCR reaction, we did not observe equal cluster densities (Figure 2).

**Figure 2.**
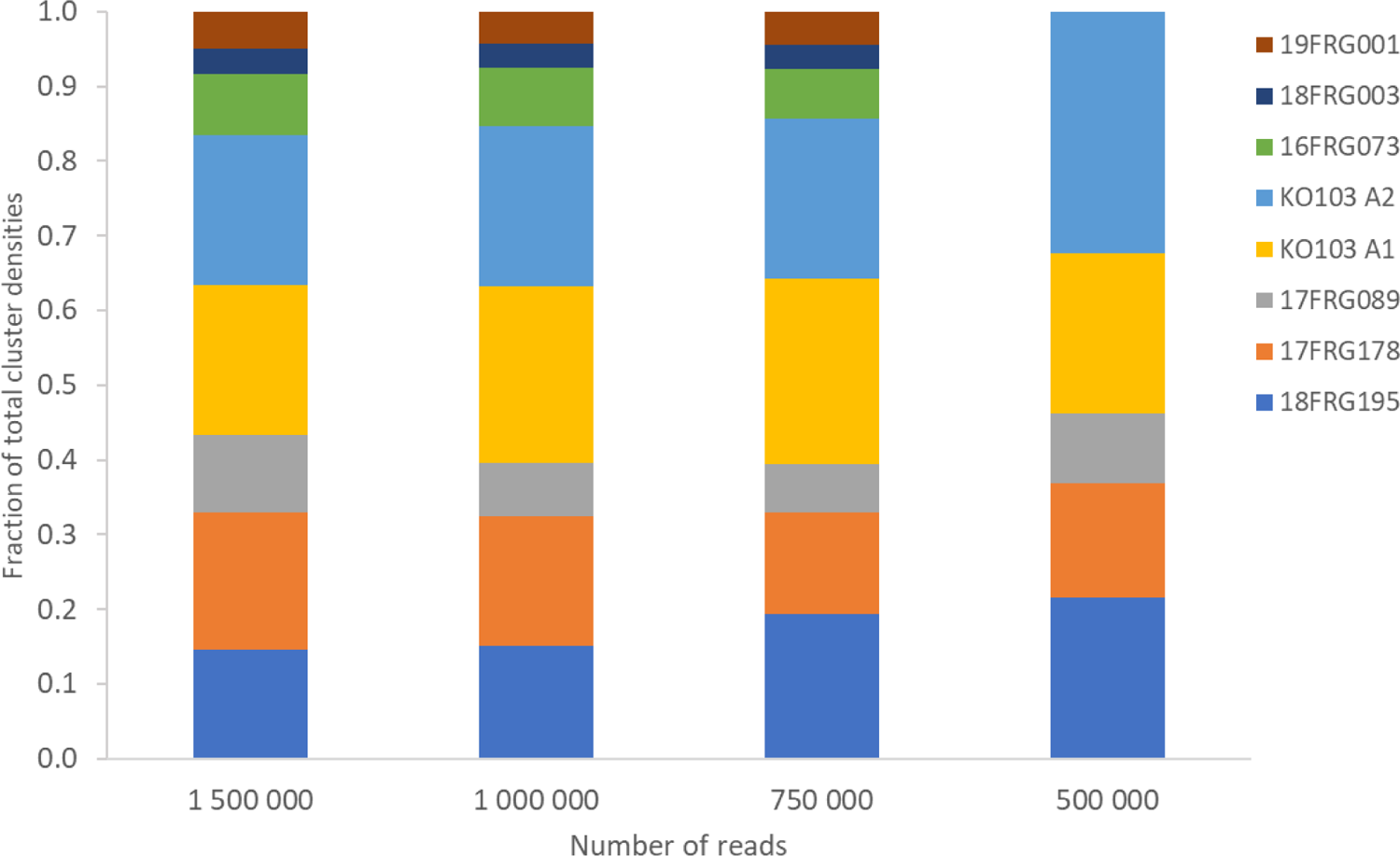
Cluster densities of each *Cyp51A* allele as a fraction of the total for 1 500 000, 1 000 000, 750 000 and 500 000 randomly sampled reads inputted into the longread UMI bioinformatic pipeline.

**Table 3.** Reads verses variants. Number of subsampled reads verses depth of clusters with 100% identity to the respective reference *Cyp51* sequences for each variant in a mock community.

| Number of reads | Cluster depth for each *Cyp51A* variant |  |  |  |  |  |  |  |
| --- | --- | --- | --- | --- | --- | --- | --- | --- |
|  | 18FRG195 | 17FRG178 | 17FRG089 | KO103 A1 | KO103 A2 | 16FRG073 | 18FRG003 | 19FRG001 |
| 1 500 000 | 1465 | 1854 | 1037 | 2030 | 2008 | 830 | 342 | 494 |
| 1 000 000 | 395 | 450 | 189 | 615 | 560 | 201 | 84 | 113 |
| 750 000 | 135 | 95 | 45 | 173 | 150 | 47 | 22 | 31 |
| 500 000 | 14 | 10 | 6 | 14 | 21 | 0 | 0 | 0 |
| 250 000 | 0 | 0 | 0 | 0 | 0 | 0 | 0 | 0 |

Variants with cluster depth of >100 were 100% identical to the reference sequences and differentiated between a one base pair shift in the promoter insert (Supplemental Figure S2). However, with increasing sequence input we noticed a concomitant increase in spurious clusters with either deletions or base changes (Table 4). All spurious clusters analysed in this study contained errors in the *Cyp51A* coding sequence, with the majority consisting of base pair deletions in a heteropolymer a/g di-nucleotide region (Figure 3). Even though the number of spurious clusters increased with the number of input sequences, the depth of the spurious clusters was approximately an order of magnitude less than the variants which were 100% identical to the reference sequences, thus easily discarded (Table 3, Table 4). We determined that approximately 100K reads per haplotype was optimal for resolution of all haplotypes and reduction in spurious clusters, while maximizing the number of samples that can be run on each flow cell.

**Figure 3.**
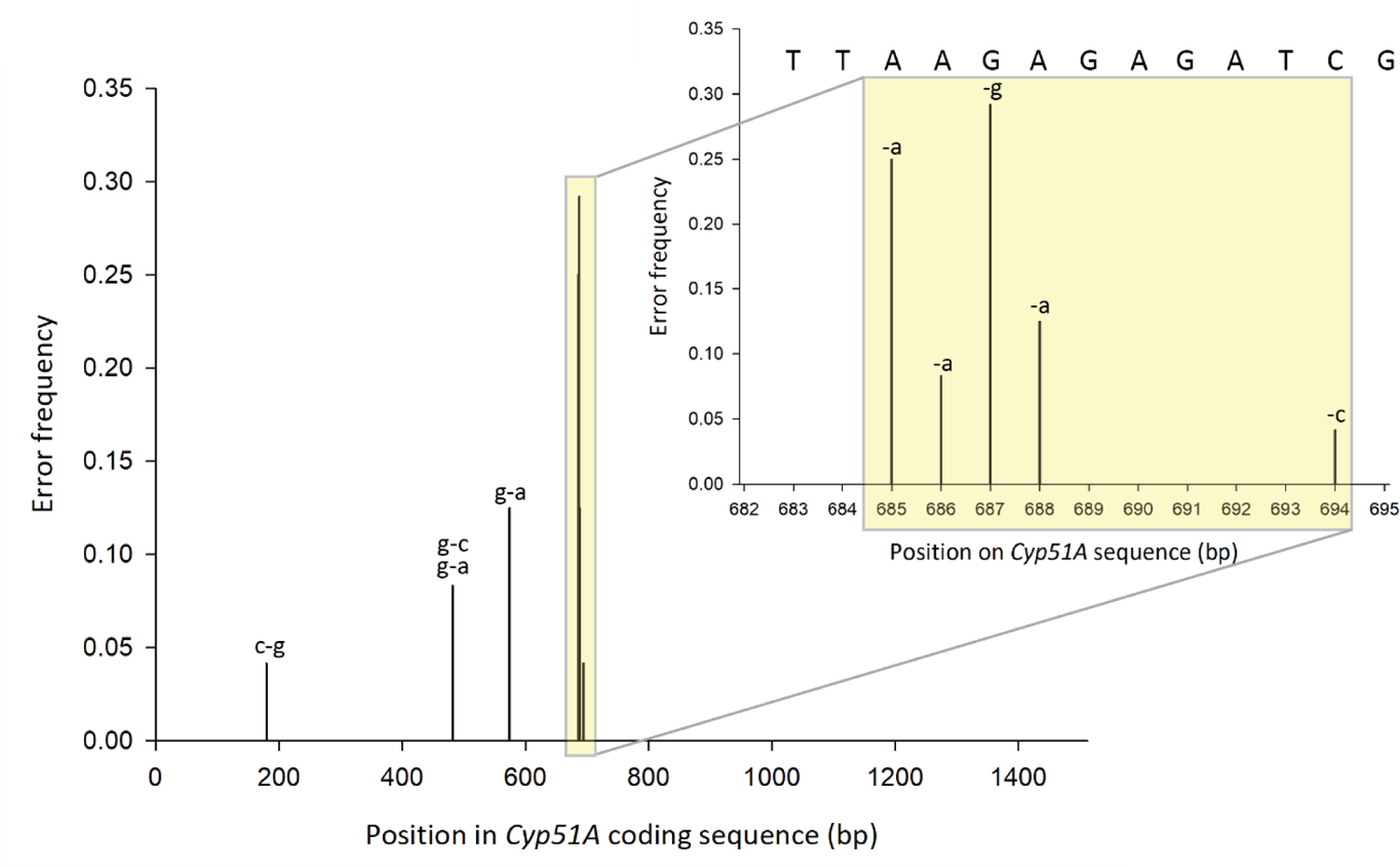
Frequency, type, and position in *Cyp51A* coding sequence of errors present in all spurious clusters observed in this study.

**Table 4.** Number of spurious clusters and depth with <100% identity to respective reference *Cyp51* sequences for each variant for mock community.

| Number of reads | Number spurious clusters for each *Cyp51A* variant (Cluster depth) |  |  |  |  |  |  |  |
| --- | --- | --- | --- | --- | --- | --- | --- | --- |
|  | 18FRG195 | 17FRG178 | 17FRG089 | KO103 A1 | KO103 A2 | 16FRG073 | 18FRG003 | 19FRG001 |
| 1 500 000 | 2 (5,5) | 2 (3,5) | 1 (4) | 1 (5) | 2 (6,5) | 1 (7) | 0 | 0 |
| 1 000 000 | 0 | 0 | 1 (4) | 2 (3,3) | 1 (4) | 0 | 0 | 0 |
| 750 000 | 0 | 1 (3) | 0 | 0 | 0 | 0 | 0 | 0 |
| 500 000 | 0 | 0 | 0 | 0 | 0 | 0 | 0 | 0 |
| 250 000 | 0 | 0 | 0 | 0 | 0 | 0 | 0 | 0 |

To validate the mutations in the *Cyp51A* mock community, we analysed the input DNA used for ONT sequencing with qPCR assays specifically designed to detect the genetic changes in each haplotype (Knight *et al*., 2023; Figure 1, Table 2) with the pure genomic DNA from each isolate as positive controls (Supplemental Figure S1). Negative controls consisted of genomic DNA from an isolate known to not contain the mutation being tested. The qPCR analysis confirmed the presence of each expected mutation associated with fungicide resistance in the seven isolates tested (Supplemental Figure S1).

### Custom workflow identifies *Cyp51* haplotype associated with azole resistance from field samples

To avoid lengthy isolation procedures to generate single spore isolates, we developed a customized workflow designed for complex cultures from leaf samples (Figure 4). We analysed five barley leaf samples from South Australia with net form of net blotch symptoms. Since the leaf samples were infected with *Ptt* (confirmed by qPCR, data not shown), two clusters representing *Cyp51A1* and *Cyp51A2* were expected, as there are two *Cyp51A* alleles in *Ptt* (Mair et al., 2016). The resulting total of 845 625 passed sequences as well as 750K, 500K, 375K, 300K, 250K and 100K randomly sampled sequences were inputted into the longread UMI pipeline. No variants resulted from either the 100K or 250K read inputs, however 300K reads and above successfully generated variants (Table 5). Like the mock community analysis, spurious clusters increased with increasing read inputs (Table 4, Table 5). Error profiles of spurious clusters were like those found in the mock community analyses (Figure 3) except for a t to c substitution in a homopolymer region eight nucleotides before the start codon in one spurious cluster (Supplemental Figure S3), but these were of discernibly low depth and thus easily discarded (Table 5). In all successful pipeline runs, two high-depth clusters resulted with two non-synonymous and one synonymous mutation in the coding region and two promoter SNPs (Supplemental Figure S3). We re-evaluated the reads/haplotype to be approximately 100K to 150K reads/haplotype for haplotype detection and minimization of spurious clusters (Table 5). The two non-synonymous coding-region SNPs caused I133V and F489L amino acid substitutions in one of the clusters, the latter associated with *P. teres* azole resistance. The c1465t substitution causing the F489L mutation was validated as the only mutation present in the five samples by qPCR (data not shown).

**Figure 4.**
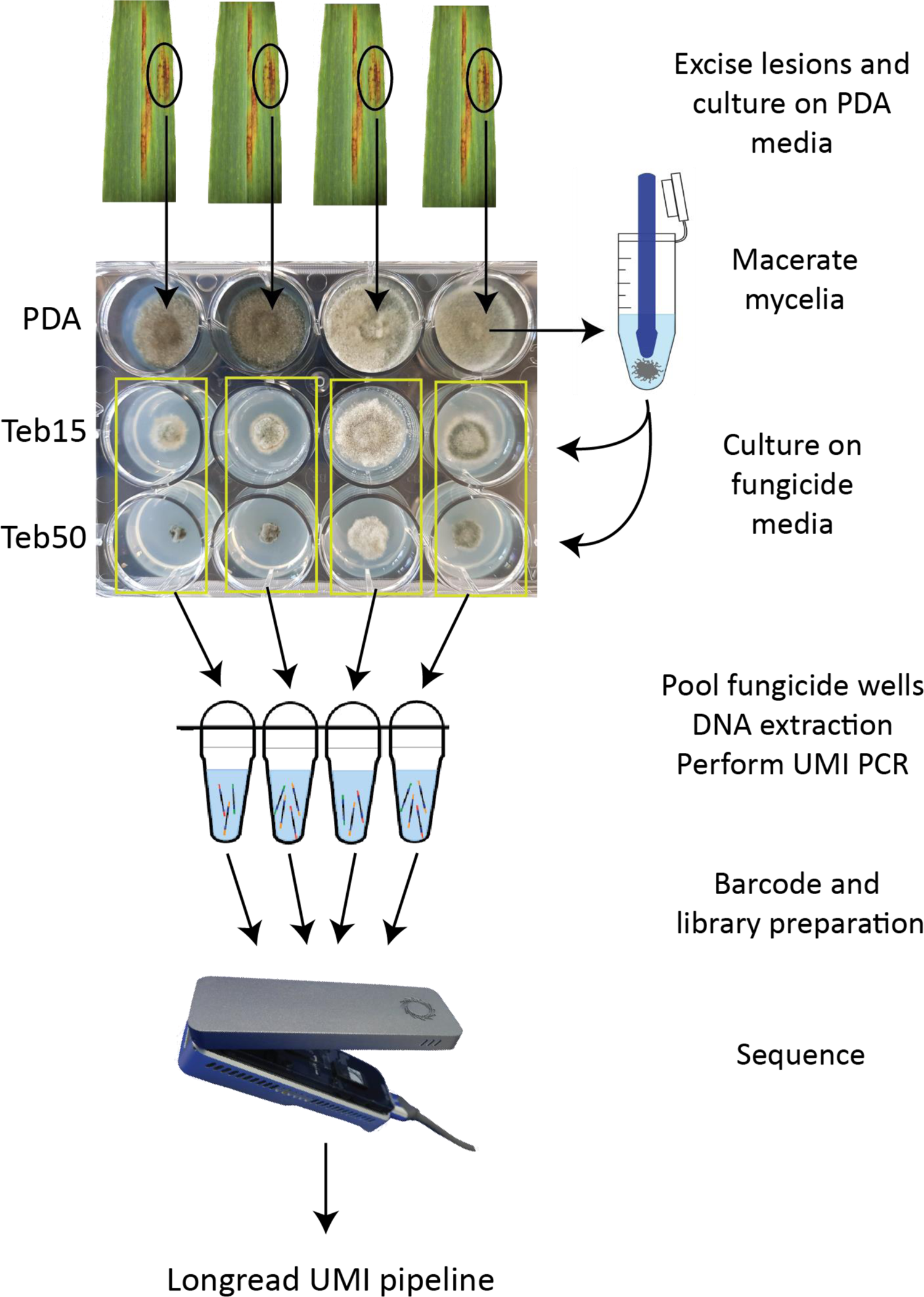
Workflow for culturing, amplification, sequencing, and bioinformatic analysis of mycelia cultured on tebuconazole amended media from infected leaf samples. Teb15 = 15 µg mL^-1^; Teb50 = 50 µg mL^-1^.

**Table 5.**
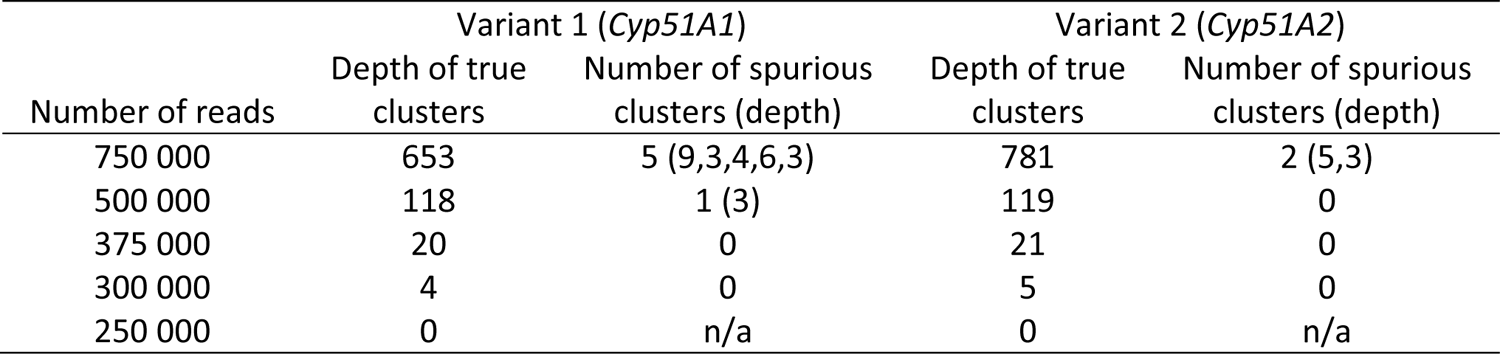
Number of subsampled reads verses true cluster depth and number of spurious clusters and associated cluster depth for each variant (*Cyp51A1* and *Cyp51A2*)

## Discussion

In this study we used the MinION, a portable hand-held sequencer from Oxford Nanopore Technologies, to sequence the azole target gene *Cyp51* from characterized mock communities and infected leaf samples from paddocks in South Australia. As a proof of concept, we amplified the promoter and coding region of *Cyp51A* from seven isolates representing a diverse array of haplotypes including promoter insertions and coding sequence SNPs from *Ptt*, *Ptm* and a hybrid. The long read UMI pipeline developed by Karst et al., 2021 resolved all insertions and SNPs of all variants with 100% identity to the respective reference *Cyp51A* sequences. Our method also discriminates between *Ptt* and *Ptm* as the nucleotide at position 1488 in the *Cyp51A* gene is a cytosine in *Ptm* and a guanine in *Ptt* in all isolates we have sequenced to date. Thus, with one sequencing assay it was possible to determine the *P. teres* form(s) present and associated azole fungicide resistance status.

Fungicide resistance is a global issue and rapid diagnosis is critical to prevent epidemic emergence. However, detection of agrochemical resistance can be complex and time consuming because one must monitor for multiple mutations in target genes. For example, in two closely related forms of *P. teres* and a hybrid the target gene for azole fungicides, *Cyp51A*, can contain promoter insertions at several locations with or without the F489L mutation, which itself can result from three different codon changes (Mair et al., 2016, Mair et al., 2020, Turo et al., 2021). Eight inserts that differ in position or sequence have been detected in the *Cyp51A* promoter of *Ptm* to date, thus it is possible that there are additional insertion sites that have not yet been characterised (Mair et al., 2020, Knight et al., 2023). Both the promoter insert and the coding sequence mutation in the same allele result in high levels of resistance (Mair et al., 2020). In the wheat pathogen *Z. tritici*, numerous coding sequence mutations and promoter modifications have also been associated with resistance to azole fungicides (Huf *et al*., 2018). This complex resistance landscape dictates the need to profile the entire target gene and promoter region to resolve the haplotype present, not individual mutations in isolation. This is especially important when profiling biologically complex field samples.

We applied our pipeline to infected barley leaf samples and used a culturing method on fungicide media developed by Knight et al. (2023). This method has three main advantages: i) bulking of fungal mycelia for PCR, ii) removal of plant DNA, and iii) selection of only fungicide resistant isolates. As expected, we recovered two high depth clusters for each field sample (*Cyp51A1* and *Cyp51A2*), one of which had two non-synonymous SNPs, I133V and F489L, the latter correlates with decreased fungicide sensitivity, while the former does not (Mair et al., 2016). The codon which gave rise to the F489L mutation was c1465t which was originally reported in *Ptm* (Mair et al., 2020) but only recently in *Ptt* (Knight et al., 2023). If this codon change had not been discovered prior, the result of an allele-specific assay would have been a false negative, highlighting the utility of our assay for comprehensive resistance profiling. Since our assay provides full sequence information on both promoter and coding sequence mutations, it can decern that F489L and one other coding sequence mutation (I133V) were present, and no promoter insertions were detected to potentially account for the growth in media amended with tebuconazole. To reach this conclusion using traditional methods would involve lengthy single spore culturing procedures and Sanger sequencing, which demonstrates the utility of our assay to deliver several critical pieces of information at once which saves time and cost.

Our assay offers distinct advantages over detection methods such as LAMP and qPCR, especially in field samples where several mutations can co-occur in one allele and several alleles likely are present in one sample. This is especially salient for pathogens without existing target gene mutation profiles, as individual mutation detection assays do not need to be developed. Also, fungal genomes are relatively large compared to viral or bacterial genomes and can be prohibitively expensive to sequence for diagnostic purposes. Amplicon sequencing targets only those genes of interest, therefore reducing the amount of data that needs to be collected and therefore cost of the assay. However, the target gene sequence must be known and if resistance is not solely due to target gene changes it will still be missed.

Recently, ONT sequencing was used to profile *Cyp51* mutations in *Z*. *tritici* mixed isolate mock communities and infected wheat leaves (Gutierrez Vazquez et al., 2022). To mitigate sequencing errors, the authors employed a filtering and correcting approach using Filtlong and Canu, respectively. The filtering step was used to increase the overall quality of the basecalls to the highest quality 10 million bases, however this resulted in substantial read loss when sequenced with the R9.4 MinION flow cell for a Phred score of 13.8 when sequencing *Cyp51* mock communities. It is important to highlight that these issues will likely decrease with increasing accuracy of the R10.4/Kit 14 flow cells and associated chemistry, which has enabled near perfect bacterial genomes without short read or reference polishing (Sereika *et al*., 2022). Nevertheless, Gutierrez Vazquez et al. (2022) obtained a per position consensus of 98 to 99% after read correction and identified a total of 16 unique haplotypes of *Z*. *tritici* across the 16 wheat leaves sampled (Gutierrez Vazquez et al., 2022). The use of UMIs for read correction differs from this approach in several respects. Consensus is determined at the individual molecule level which results in a very low error rate (mean error rate = 0.0042%) and since two UMIs are required for binning, the chimera rate (<0.02%) is also very low (Karst et al., 2021). When using the ONT R10.3 flow cell to sequence microbial mock communities, the chimera filtering function of the UMI pipeline removed 4.8% of chimeric UMI bins (Karst et al., 2021), highlighting the ability of this pipeline to eliminate these potentially erroneous results. Low error rate and chimera filtering are two important considerations when accurately detecting particularly low abundance variants in complex samples. The high depth clusters generated in this study were 100% identical to the reference sequences and no chimeras were detected. Although we did detect approximately 100X lower depth clusters which were not 100% identical, these errors were mostly in homopolymer regions, which is a feature of ONT sequencing and were easily discarded.

Amplicon sequencing was also used to sequence the target genes of azole, SDHI and QoI fungicides along with several genes used for phylogenetic classification in the phytopathogen *Z. tritici* (Samils et al., 2021) using PacBio sequencing. PacBio sequencing also suffers from relative high error rates of around 13% (Karst et al., 2021), however the Sequel platform (Pacific Biosciences, Menlo Park CA, United States) overcomes this by utilizing circular consensus sequencing which resulted in a Phred quality score above Q20 or 99% accuracy (Samils et al., 2021). Single spored isolates were generated and tagged using a two-step PCR approach where the first PCR introduced a ‘heel’ sequence which was then tagged in the second PCR to enable differentiation of each isolate in downstream data analysis (Samils et al., 2021). The amount and diversity of data generated in this assay is commendable and is useful for generating both resistance and phylogenetic profiles of isolates in large populations. Our approach differs in that we used UMIs to tag and differentiate individual molecules in complex mixed samples that contain more than one isolate, therefore eliminating the need for isolation of individual strains. Thus, we can detect the full suite of resistance haplotypes present in a lesion without the need for isolations and with SNP-level accuracy due to the generation of molecule-specific consensus sequences.

Our method also exploits the portability of the MinION sequencer, which has the potential to enable diagnostics in the paddock. The MARPLE disease diagnostic and surveillance tool has been recently reported to identify and assign strains of *Puccinia striiformis* f. sp. *tritici* to distinct lineages with a single sequencing assay using the MinION (Radhakrishnan *et al*., 2019). Similar to the current study, genetically distinct regions were identified and selectively amplified. The authors comment that the addition of fungicide target genes such as *Cyp51* and the four succinate dehydrogenase subunits would be useful in real time monitoring of known mutations in these genes. It is important to accurately characterize SNP level changes to discover new uncharacterized mutations which would require very high accuracy sequencing techniques. The UMI approach outlined in this study offers higher accuracy and lower chimera rates than previously possible on long read sequencing platforms such as the MinION (Karst et al., 2021).

We found systemic errors in clusters occurring around single and di-nucleotide repeats in the *Cyp51* gene. These systemic homopolymer-associated errors are a feature of ONT sequencing and were also identified in standard microbial community profiling using UMIs and the ONT platform, mostly as deletions in long cytosine and guanine homopolymers and guanine insertions in non-homopolymer regions (Karst et al., 2021, Delahaye & Nicolas, 2021). Although this phenomenon did result in more clusters than were expected, this was not deemed to be problematic as the clusters could easily be discarded due to their relatively small (approximately 100X lower) cluster depth.

Another important factor in the success of the UMI approach is input DNA amount which must be optimised to achieve the desired single molecule coverage and sequencing yield required (Karst et al., 2021). We determined the optimal amount of input DNA to be 1.3 ng, but also empirically tested amounts lower and higher than this and found DNA amounts less than 1.3 ng did not yield product while amounts higher than 1.3 ng did not yield sufficient cluster depth for the bioinformatics pipeline to function. If this method is to be applied directly to infected leaf samples, an intermediate qPCR step would likely be required to determine the amount of pathogen DNA present in the leaf sample, as we found this to be a critical factor to the success of this method.

The DNA used as input for the first PCR reaction in the *Cyp51A* mock community consisted of an equimolar amount of each of the seven *P*. *teres* isolates, we did not observe an equal cluster density for each of the true variants. Karst et al. (2021) also observed a discrepancy between the theoretical relative abundance of the rRNA operons from the mock community supplied by the manufacturer and the observed relative abundances when sequenced using the UMI pipeline. The authors hypothesized this could be due to biased fragments size, difference in growth rates and primer mismatches are possible reasons. We do not expect these issues to be relevant here as equimolar amounts of genomic DNA were added to the PCR reaction, the amplicons are of similar size and there should theoretically be no primer mismatches among the targets. It is interesting to note that the cluster densities for KO103A1 and KO103A2 were nearly identical, which is expected since the two alleles are from the same isolate. It is possible that the observed differences are isolate dependent. Compounds such as oxidized polyphenols and polysaccharides are known to interfere with PCR (Inglis *et al*., 2018) and genomic DNA isolations from *P*. *teres* have been reported to be viscous and discoloured, which required a partial digestion of the cell wall to obtain high quality DNA for PacBio sequencing (Syme *et al*., 2018). We have anecdotally observed differences in pigmentation and viscosity of DNA extractions from different isolates of *P. teres*, and this could potentially cause differential amplification when they are mixed.

While the UMI sequencing approach allows entire gene variants to be described, it currently lacks quantification of allele frequencies. This suggests that quantitative platforms such as qPCR or digital PCR are still complementary for the detection process, especially as they are likely more sensitive and able to detect fungicide resistance mutations as low as 0.2% (Zulak et al., 2018). Also, our current field sample pipeline relies on a lengthy culturing step for selective growth of resistant isolates, similar to detection pipelines using qPCR or digital PCR (Knight et al., 2023). Ideally improvements will be devised which would allow DNA to be extracted directly from diseased leaves for UMI sequencing and amplicon diversity assessments.

In this study, we adapted a UMI PCR-based sequencing method to characterize SNP-level mutations in the azole fungicide target gene *Cyp51*, illustrating the broad applicability of this approach. To improve the sample to result timeline, we are currently trialling direct *Cyp51* SNP detection from infected leaf samples which would eliminate the culturing step. It is also expected that the amount of data required for accurate consensus sequence generation and homopolymer associated errors will decrease with increasing accuracy of ONT sequencing (Sereika et al., 2022). These improvements would result in a decrease in the per sample sequencing cost, as more samples could be analysed per flow cell. Our approach could also be expanded to include phylogenetically relevant genes for simultaneous lineage profiling as highlighted in previous studies, although this would reduce the data yield for fungicide target genes and therefore the number of samples that can be included in a single run. This method could also be applied to other agrochemical resistance systems such as herbicides, insecticides and bactericides where accurate and rapid SNP level detection of new mutations is critical.

## Acknowledgements

The authors would like to thank Johannes Debler for help with GPU base calling. This study was supported by the Centre for Crop and Disease Management, a joint initiative of Curtin University, and the Grains Research and Development Corporation – research grant CUR00023.

## Supplementary Figure and Tables

**Supplementary Figure S1.**
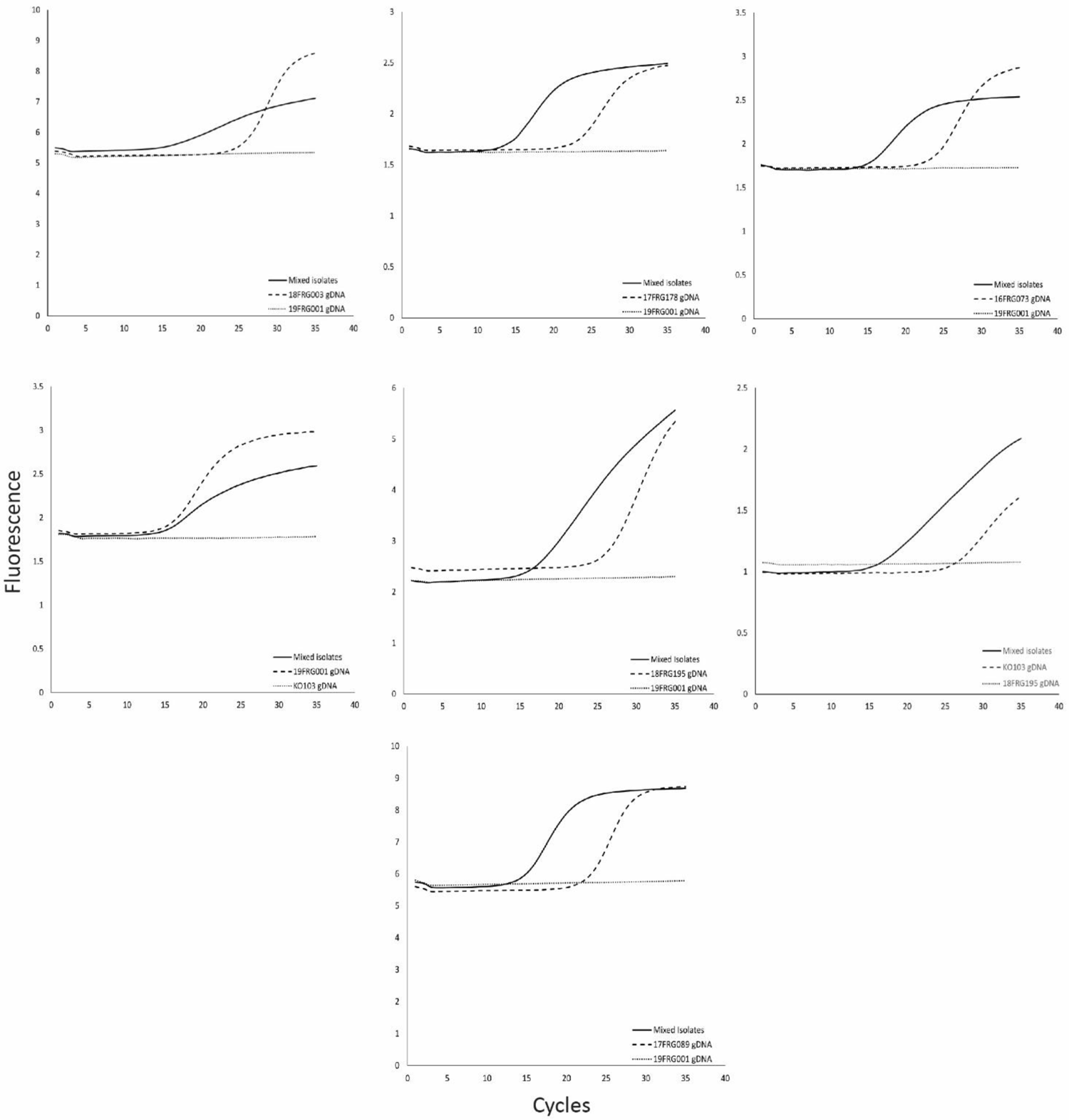
Quantitative PCR validation of UMI PCR input DNA for sequencing of *Cyp51A* mock community. Positive controls consisted of genomic DNA from an isolate known to contain the mutation being tested and are shown as a dashed line. Negative controls consisted of genomic DNA from an isolate known to not contain the mutation being tested and are shown as a dotted line.

**Supplementary Figure S2.**
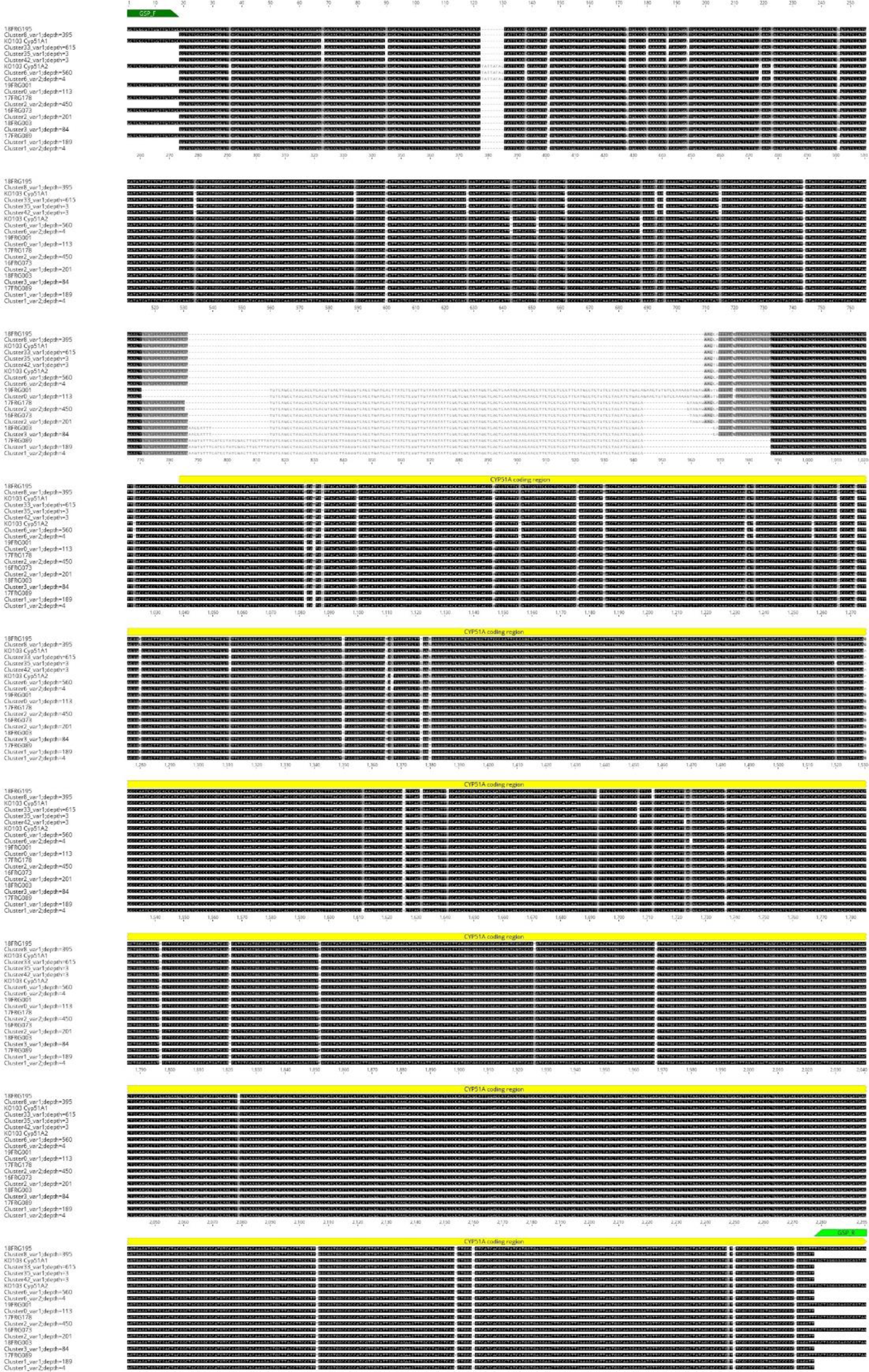
Nucleic acid alignment of clusters generated from UMI PCR of *Cyp51A* genes from seven isolates of *P. teres* compared to reference sequences when inputting 1 000 000 reads. GSP_F and GSP_R = gene specific primers used for UMI PCR (Table 2). Shading is as follows: black = 100% similar, dark grey = 80-99% similar, medium grey = 60-79% similar, light grey = less than 60% similar. Image created using Geneious 2022.1 created by Biomatters.

**Supplementary Figure S3.**
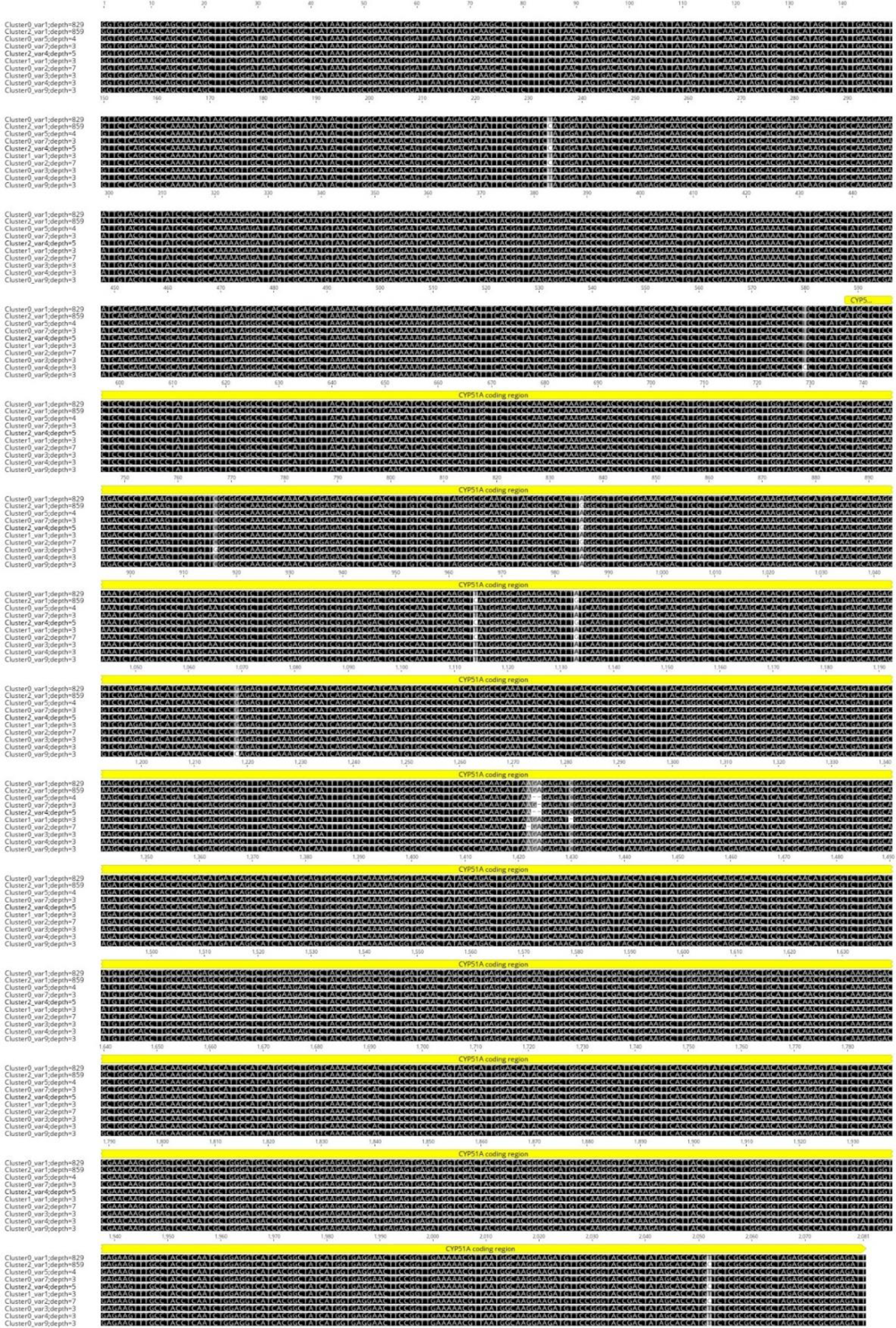
Alignment of clusters from field samples. Nucleic acid alignment of clusters generated from UMI PCR of *Cyp51A* genes from five leaf samples infected with *Pyrenophora teres* f. *teres* when inputting all reads. Shading is as follows: black = 100% similar, dark grey = 80-99% similar, medium grey = 60-79% similar, light grey = less than 60% similar. Image created using Geneious 2022.1 created by Biomatters.

